# Rest is Required to Learn an Appetitively-Reinforced Operant Task in *Drosophila*

**DOI:** 10.1101/2020.08.28.272047

**Authors:** Timothy D. Wiggin, Yung-Yi Hsiao, Jeffrey B. Liu, Robert Huber, Leslie C. Griffith

## Abstract

Maladaptive operant conditioning contributes to development of neuropsychiatric disorders. Candidate genes have been identified that contribute to this maladaptive plasticity, but the neural basis of operant conditioning in genetic model organisms remains poorly understood. The fruit fly *Drosophila melanogaster* is a versatile genetic model organism that readily forms operant associations with punishment stimuli. However, operant conditioning with a food reward has not been demonstrated in flies, limiting the types of neural circuits that can be studied. Here we present the first sucrose-reinforced operant conditioning paradigm for flies. In the paradigm, flies walk along a Y-shaped track with reward locations at the terminus of each hallway. When flies turn in the reinforced direction at the center of the track, sucrose is presented at the end of the hallway. Only flies that rest early in training learn the reward contingency normally. Flies rewarded independently of their behavior do not form a learned association but have the same amount of rest as trained flies, showing that rest is not driven by learning. Optogenetically-induced sleep does not promote learning, indicating that sleep itself is not sufficient for learning the operant task. We validated the sensitivity of this assay to detect the effect of genetic manipulations by testing the classic learning mutant *dunce*. *Dunce* flies are learning impaired in the Y-Track task, indicating a likely role for cAMP in the operant coincidence detector. This novel training paradigm will provide valuable insight into the molecular mechanisms of disease and the link between sleep and learning.

## INTRODUCTION

Learning is a broadly conserved, highly regulated, and health relevant function of the nervous system. Learning updates the frequency of behaviors to reflect stimulus predictability in an animal’s environment. The associative forms of learning transfer the value of an innately valued stimulus (an unconditioned stimulus or US) to an associated predictor, either a behavior or cue (Fanselow and Wassum, 2015). US association with internally-generated behavior (e.g. locomotion, static posture, lever press) produces “operant conditioning” across a wide range of animal species (Skinner, 1948; Kimble et al., 1955; Susswein et al., 1986). Operant conditioning allows the animal to modify its behavior to increase the likelihood of obtaining rewarding stimuli and decrease the likelihood of encountering aversive stimuli.

Operant conditioning to reward or relief from punishment incorporates a positive feedback loop – learning increases the generation of the behavior, which in turn increases reward frequency, which strengthens the learned association. This type of positive feedback loop is hypothesized to contribute to diverse neuropsychiatric disorders including childhood anxiety, compulsive behaviors, and chronic pain (Ollendick et al., 2001; Korff and Harvey, 2006; Chóliz, 2010; Gatzounis et al., 2012). Genome-wide association studies have identified candidate genes that increase susceptibility to these operant conditioning-associated disorders (Smith et al., 2016; Levey et al., 2020; Smit et al., 2020). However, despite the relevance to human health, the neural basis of operant conditioning in genetic model organisms remains incompletely understood. It is not currently possible to trace a neural circuit of operant conditioning in animals more complex than *Aplysia californica* (Nargeot and Simmers, 2011), nor has there been a genetic screen for molecular components of operant learning in model organisms. A promising system to address this gap in knowledge is the fruit fly *Drosophila melanogaster*. Much of the known molecular machinery underlying learning and memory was first discovered using genetics in the fly and these molecules have subsequently been shown to be essentially identical in humans (Greenspan and Dierick, 2004). Furthermore, a draft map of the neural connections in a fruit fly hemi-brain has been recently published which, along with advanced genetic tools, greatly facilitates mapping complex neural circuits (Pfeiffer et al., 2010; Xu et al., 2020).

Operant conditioning has been studied extensively in flies, but only limited progress has been made in understanding circuit-level mechanisms. There have been many operant conditioning paradigms reported in flies: geotaxis training (Murphey, 1967), leg position conditioning (Booker and Quinn, 1981), proboscis extension suppression (DeJianne et al., 1985), flight simulator heat avoidance (Wolf and Heisenberg, 1991), conditioned place preference (Wustmann et al., 1996), social freezing (Kamyshev et al., 1997), and left-right navigation in tethered ball-walking (Nuwal et al., 2012). However, a pair of landmark publications (Brembs and Plendl, 2008; Brembs, 2009) demonstrated that when predictive sensory cues are available, flies preferentially learn these sensory cues and block the formation of operant conditioning. This finding dramatically compromises a number of paradigms that intended to test operant learning in flies, since predictive sensory information present during training may have inhibited operant learning. The remaining purely operant learning paradigms that are routinely used in flies, flight simulator heat avoidance and proboscis extension suppression, have two important limitations. First, they use restrained fly preparations, which unavoidably alter animal behavior (Stowers et al., 2017). Second, they use an aversive US which may not recruit the full repertoire of US pathway neurons (Liu et al., 2012) and may use neurons outside the brain for learning (Booker and Quinn, 1981).

In order to extend the range of operant conditioning paradigms in flies, we developed a positively reinforced, self-paced, operant training task for untethered flies, which we call the Y-Track. Surprisingly, we found that this operant training paradigm only produces a major change in behavioral frequency in a relatively small subset of experimental animals that rest during training. This surprising finding further reinforces the importance of rest for learning (Maquet, 2001) and opens a new avenue for measuring this link in a single-session paradigm.

## MATERIALS AND METHODS

### Experimental Animals

Flies were raised on cornmeal-dextrose-yeast food in bottles at room temperature or in a 25 C incubator with a 12 hour:12 hour light:dark cycle. Wild type (WT) flies were from the Canton-Special (CS) background. Transgenic flies were obtained from the Bloomington Drosophila Stock Center (BDSC) and Vienna Drosophila Resource Center (VRDC) as follows: *P{VT058968-GAL4}attP2* (VT058968-GAL4, VDRC# 204550), *P{w[*+*mW.hs]=GawB}104y* (104y-Gal4, BDSC# 81014), *PBac{y[*+*mDint2] w[*+*mC]=UAS-ChR2.XXL}VK00018* (UAS-ChR2.XXL, BDSC# 58374), and *dnc^1^* (BDSC# 6020). Flies with Gal4 and UAS transgene insertions were outcrossed to a CS background for several generations because we found that *white* knock-out backgrounds may be learning deficient in this task (*data not shown*).

### Design of the Y-Track Apparatus

The Y-Track conditioning apparatus was designed as a 4 layered structure (Figure 1). The first (top) layer of the structure was a 3D printed holder for a USB camera (ELP-USBFHD01M) and 3.6mm S-mount lens facing downward toward the track. The second layer was a mount for a red filter (Tiffen #25 Red) to block light blue and green light from optogenetic activation and light landmark experiments. These top layers are supported by four 3D pillars on each side of the apparatus. Red LEDs (630nm, Vishay VLDS1235G) were attached to each pillar and illuminated the Y-Track area. The third layer of the apparatus was the Y-Track itself. Two versions of the Y-Track were tested: a square-walled track and a curved floor track. In the square-walled track, the width of the hallways was 3.5 mm and the height of the hallways was 2.5 mm. In the curved floor track, the track surface was described by a circular arc with a diameter of 9 mm. The width of the top of the hallways was 6.7 mm and the height at the middle of the hallways was 1.5 mm. In both tracks, each of the three hallways that made up the Y shape was 20mm long, the hallways met in the middle of the track, and the hallways had 120° radial spacing. At the end of each arm of the maze was a circular plastic holder for reward filter paper (“food circle”) securely screwed to a servomotor (Towerpro MG91). Each food circle had two filter paper slots, one for a sucrose-soaked filter paper and the other for a water-soaked filter paper. The top of the Y-Track was covered by a clear acrylic plate with a small hole for aspirating flies, machined by the Brandeis University Machine Shop. A small 3D frame with 3 RBG LEDs (Broadcom HSMF-C114) was superglued below the Y-Track to deliver optogenetic stimulation and landmark location cues. The fourth (bottom) layer of the Y-Track apparatus is a frame that positions the servomotors correctly relative to the Y-Track and secures the entire apparatus to the base. Modelling of the 3D printed components was done using Autodesk Fusion 360 (San Rafael, CA). These components were fabricated from Polylactic Acid (PLA) filament in the Brandeis MakerLab.

**Figure 1.**
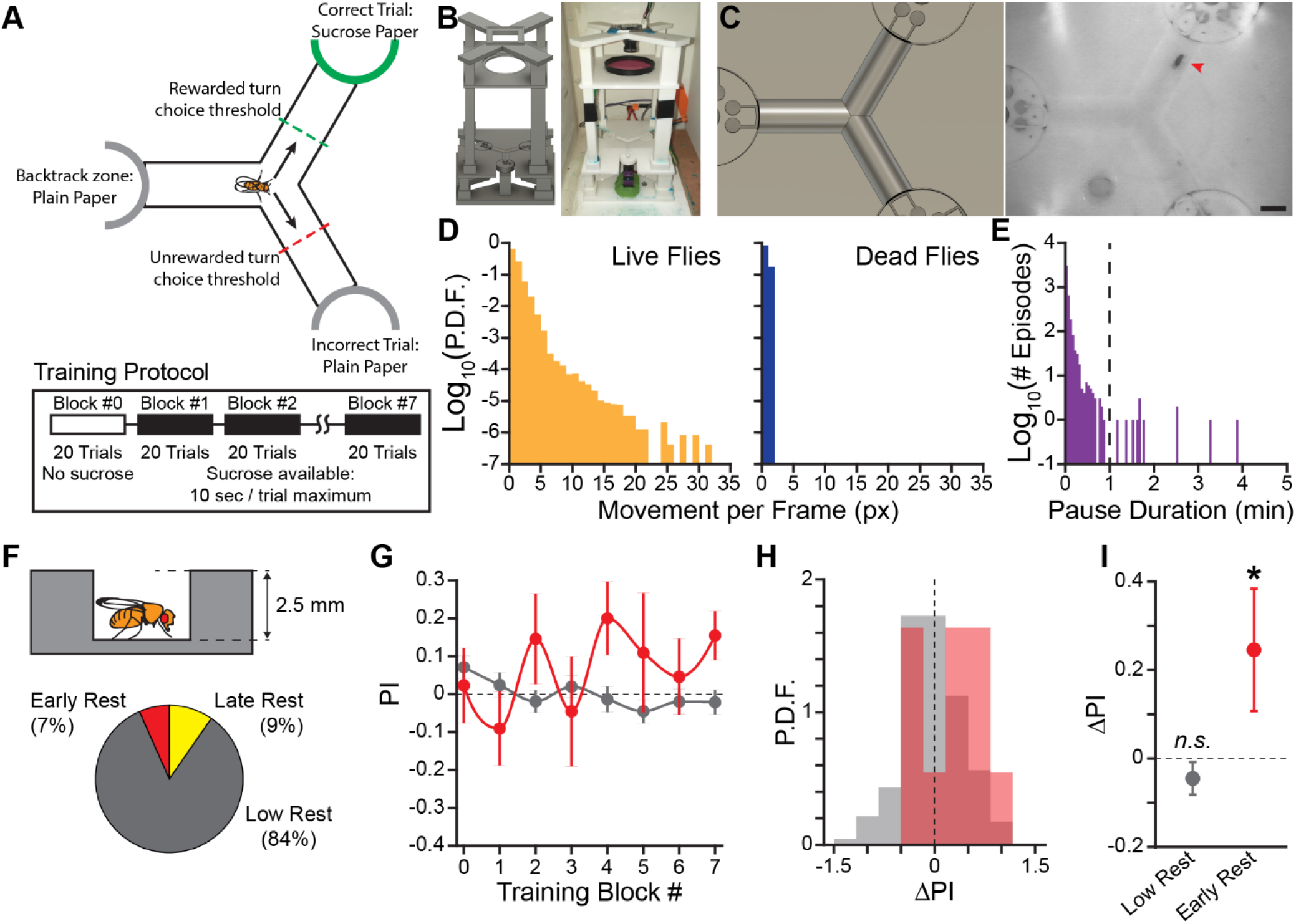
Flies That Rest Learn the Operant Contingency in a Novel Sucrose Seeking Task. (A) Diagram of operant conditioning paradigm. Sucrose is presented at the end of the track in the reinforced direction. A trial is completed after the fly crosses the choice threshold leaving the center of the track. Each training block is 20 trials, reward presentation begins in training block #1. Sucrose is made available for 10 seconds following the fly crossing the choice threshold. (B) Rendering of the 3D model of the novel apparatus used for 3D printing (*Left*) and a photo of a fully assembled apparatus (*Right*). (C) Field of view of overhead camera in the 3D render (*Left*) and a video frame of a fly navigating the Y-Track (*Right*). Red arrowhead indicates the position of the fly. Scale bar: 5 mm. (D) Probability Distribution Function (P.D.F.) histogram of movement (in pixels) per frame (approx. 50ms) for live flies (*Left*) and dead flies (*Right*). One pixel of movement is approximately 90 μm. (E) Histogram of locomotor pause durations from 12 hour recordings of locomotor behavior (*n* = *2* flies, 4127 episodes). Dashed line indicates the 1 minute threshold to distinguish short pauses from rest episodes. (F) Schematic of hallway geometry with representative fly for scale (*Top*), Fraction of flies with each rest phenotype (*Bottom*). (G) Mean turn direction Preference Index (PI) for each training block. Points plotted in color are flies with early rest (drowsy), points plotted in grey are flies with low rest (restless). (H) Probability Distribution Function (P.D.F.) histograms of change in turn preference index (ΔPI) between training block #1 and #7. Drowsy flies are plotted in red, restless flies are plotted in grey. (I) Mean ΔPI of low rest and early rest flies. Error bars are standard error of the mean, *n.s*.: groups are not significantly greater than zero, * groups are significantly greater than zero.

This apparatus was controlled by a custom Java program running on an Udoo X86 Advanced Plus single-board computer. The JavaGrinders library was used to interface with the camera and servomotors (Donelson et al., 2012). Servomotors were controlled via a Phidget Advanced Servo (Phidgets, Calgary, CA). Red LEDs illuminating the Y-Track were powered by a BuckPuck (LuxDrive, 03021-D-E-700). The Y-Track apparatus and electronics were securely mounted inside a custom-built particle board box to provide environmental isolation. The internal walls of the box were painted white to reduce visual cues and a 120mm low-noise ventilation fan was installed to prevent overheating. Each Y-Track single-board computer was connected to a central control computer and controlled remotely via Virtual Network Computing (VNC). Code and 3D models are available on GitHub (https://github.com/Griffith-Lab).

### Learning Assay

Flies were collected when 0-1 days old and housed in mixed-sex vials for 24 hours to allow mating. The flies were then screened under light CO_2_ anesthesia and stored in single-sex vials of up to 20 flies each. Flies were housed in a 25 C incubator that was only accessed during the lights-on period for 7 days prior to the experiment to ensure circadian entrainment. Each vial of flies was flipped onto fresh food at 5 days old (48 hours before training), flipped onto a food-deprivation vial at 6 days old (24 hours before training), and trained in the Y-Track at 7 days old. Food-deprivation vials were made by inserting a kimwipe soaked with 1mL of tap water into an empty vial.

Prior to introducing the flies into the Y-Track, filter paper was prepared for the food circles by pipetting 30μL of 2M sucrose solution (reward stimulus) or tap water (neutral stimulus) and allowing the paper to dry overnight. The dried filter papers were securely placed into the food circles, and the positioning of the servomotors was adjusted to ensure that the flies could access only the intended stimulus and were not able to escape. Following the final positioning of the motors, a reference image of the Y-Track without a fly present was captured for image background subtraction during the experiment. Finally, the reward direction for the experiment and the sex of the experimental animal was chosen based on the experimental design. In the square-wall experiment, half of the flies were rewarded for turning right and the other half were rewarded for turning left. We found no difference between training efficacy between the reward directions, so subsequent experiments used left-turn rewards for all animals. In the square-wall experiment, half of the animals were male and half were female. We did not find a significant difference in learning between male and female flies in this experiment, but we found that males were more susceptible to starvation stress than females, consistent with previous literature (Jang and Lee, 2015). In order to reduce inter-individual variability, subsequent experiments used only female flies. Training was performed during the lights-on period of the fly’s circadian day, Zeitgeber Time (ZT) 0-9.

At the beginning of a standard training session, a single fly was aspirated out of the food deprivation vial into a Y-Track apparatus, and the lid of the track was secured in place to prevent escape. The Java control program was initialized to run the remainder of the experimental protocol, as follows: 1) The fly was given five minutes to acclimate to the maze with no sucrose presented. This acclimation time was a fixed interval and not dependent upon fly locomotion. 2) Block 0 began (Trials 1-20) and left/right turn decisions were recorded. No sucrose was presented. Block 0 was a locomotion-dependent acclimation period to ensure the fly is navigating the track. 3) Block 1 began (Trial 21). All servomotors turn to present the sucrose-soaked filter paper to the fly. A trial was initiated when the fly came within 6 mm of the center of the arena (“center zone”). If the fly back-tracked into the arm of arena it previously occupied, the servomotor turned and presented the water-soaked filter paper until the fly re-entered the center zone, but the trial continued. If the fly turned in the unrewarded direction, the servomotor turned and presented the water-soaked filter paper, ending the trial. If the fly turned in the rewarded direction, the servomotor did not turn, and the fly was given access to the sucrose-soaked filter paper, ending the trial. After a rewarded trial, the fly was given 10 seconds to consume sucrose. If the fly did not initiate a new trial by entering the center zone within 10 seconds, the servomotor turned and presented the water-soaked filter paper until the fly initiated a new trial. 4) After Trial 160, the fly was removed from the Y-Track.

In the open-loop, yoked-control experiment (Figure 3), acclimation and trials were defined exactly as in the standard protocol. However, instead of trials being rewarded or unrewarded based on turn direction, trial outcome was determined by the reward/non-reward sequence of a previously run fly. In the visual land-mark experiment (Figure 4), the training protocol was the same as the standard experiment but a single green LED (525 nm peak) was illuminated under the Y-Track. In optogenetics experiments, flies were fed food supplemented with either 1.6 mM all trans retinal (ATR) dissolved in ethanol (4% final concentration), or ethanol alone as a Vehicle control. Food deprivation vials were also supplemented with ATR or Vehicle in the same concentration as the food. During ATR supplementation flies were housed in the dark to prevent premature activation of ChR2.XXL expressing neurons. During training, blue LEDs (470 nm peak) were illuminated for 5 minutes after the initiation of Trial 50 (Block 2).

### Quantification of Activity and Rest

During training, the frame-by-frame coordinates of each fly (acquired at 20Hz), trial times, and trial outcomes were recorded. Coordinates were processed following training to remove incorrect detections, which were identified by fly coordinates outside the Y-Track region or a change in position faster than a fly could plausibly execute (Mendes et al., 2013). Gaps in data introduced by this error checking were filled by a linear interpolation of fly position. The position of the fly over time was used to determine when the fly was active: activity episodes were continuous periods of movement greater than 1 px / frame (approx. 50 ms), in which the fly also exceeded 2 px / frame at least once. The activity/inactivity sequence was used to find rest episodes, which were then used to classify flies as drowsy (3 minutes or more of rest in Block #1-3), late-resting (9 minutes or more minutes of rest in Block #4-6), or restless. Finally, the error corrected sequence of left/right turn directions was compared to the real-time turn direction determined during training. If more than 10% of the trial outcomes differed between the real-time and *post-hoc* methods, the data from the fly was excluded from further analysis due to suspected tracking errors. Analysis code implementing this process is available on GitHub (https://github.com/Griffith-Lab).

### Simulation of Turn Behavior with Turn-Direction and Alternation Bias

The interaction of turn-direction preference and alternation preference was determined using a simulation designed to be analogous to the Y-Track. Simulated, or *in silico*, flies occupied one of three positions: *P1*, *P2*, or *P3*. Like the arms of the Y-Track, these positions were arranged clockwise in a circle, so that *P1* was to the “left” of *P2*, *P2* was to the left of *P3*, and *P3* was to the left of *P1*. At each simulation step, the *in silico* fly moved from its current position to a new position. The *in silico* fly was given a pre-determined preference for/against alternation and for/against left-hand turns in the range −2 to 2, where 0 was indifference. The choice of destination was based on a biased random process based on the previous location occupied and the turn-direction preference. Alternation and turn-direction preference could combine additively or offset one another, depending on the sequence of positions visited. These preferences were combined into *net bias* = *2*^*(alternation preference* + *left-turn preference)* or *2*^*(alternation preference* - *left-turn preference)*, for the additive and offsetting case, respectively. This net bias was used to find the turn probability: *P* = *1* – *2*^*(-net bias)*. Each simulation was run for 1000 iterations to create a statistically representative sample of *in silico* behavior. Code implementing this simulation is available on GitHub (https://github.com/Griffith-Lab).

### Confocal Imaging of Fly Brains

To measure the expression pattern of Gal4 driver lines, adult fly brains were dissected at 2-5 days old and prepared for imaging. Dissection, fixation, staining, and mounting were performed according to the FlyLight protocol (Meissner et al., 2018). Primary antibodies were rabbit anti-GFP (Invitrogen, RRID:AB_221569) and mouse anti-Bruchpilot (nc82, RRID:AB_2314866) (Wagh et al., 2006). Samples were mounted in DPX (dibutyl phthalate in xylene, Sigma) and imaged with a Zeiss LSM 880 confocal microscope.

### Experimental Design and Statistical Analysis

Experiments were designed with change in Preference Index (ΔPI) as the primary measure of learning. Preference Index (PI) was defined as the preference for the rewarded turn direction and equal to (#correct turns - #incorrect turns)/#total turns. ΔPI was defined as the difference between the PI in Block 7 and Block 1. Learning was measured by performing a right-side, one-sample bootstrap test testing the null hypothesis that mean ΔPI is less than or equal to zero. The two-sample, two-sided bootstrap test was used to compare the magnitude of learning in groups of flies. Hypothesis testing was performed using 50,000 bootstrap samples; the minimum resolvable p-value using this method is approximately 0.0001. Comparisons of behavior between flies in square-walled and curved-floor Y-Tracks were performed using Two-Factor ANOVAs and Tukey-procedure protected *post-hoc* tests. Within figures, groups that are not statistically different are identified by the same letter assignment. Coordinate data and statistics were calculated using MATLAB (MathWorks, Natick, MA) and 0.05 was used as the p-value for statistical significance. The analysis code used to calculate bootstrap statistical tests is available on GitHub (https://github.com/Griffith-Lab).

## RESULTS

### Flies That Rest Learn the Operant Contingency in a Novel Sucrose-seeking Task

We used an ethology-informed approach to design a positive-valence operant conditioning paradigm. Flies locomote spontaneously while awake (Martin et al., 1999), forage for food in open fields (Hughson et al., 2018), and are adept at navigational tasks (Warren et al., 2019). We therefore used food-seeking and navigation as the central features of the learning paradigm (Figure 1A). Flies are individually loaded into a Y-shaped track (Y-Track). At the terminus of each arm of the track is a reward location that can be switched between a food reward and a neutral stimulus. Food reward is only available when the flies turn in the in the rewarded direction (*i.e*. left or right) relative to their previous location in the track. Because the rewarded choice is defined relative to the location of the animal, the location of the next rewarded location changes based on the previous behavioral choice - no single arm of the track is preferentially rewarded. Over many left/right choices (“trials”), the turn preference index (PI) is calculated for blocks of 20 trials as *PI* = *(# correct turns* – *# incorrect turns) / # total turns*. Because baseline left/right turn preference is idiosyncratic to individual flies (Buchanan et al., 2015), learning is measured as the change in PI (ΔPI) across training to determine if the flies increase their preference for turning in the direction of food reward.

Implementing this task in a physical apparatus required satisfying several design constraints (Figure 1B,C). First, the animal must be alert, healthy, and active to engage in spontaneous locomotion and learning. We included a loading port in a tightly fitted Y-Track lid that allowed us to load and remove flies without anesthesia using gentle aspiration. Second, the apparatus must include a detector element that records the performance of the reinforced behavior in real time. We used JavaGrinders real-time video tracking to measure locomotor behavior and turn choices (Donelson et al., 2012). Third, the apparatus must be able to actuate reward delivery based on the behavioral contingency. We used closed-loop control to allow the real-time tracker to activate servomotors at the terminus of each arm of the Y-Track and present either 10 seconds of access to a food reward (filter paper pre-soaked with 2M sucrose) or a neutral stimulus (plain filter paper). Importantly, the servomotors are positioned to present sucrose at both termini while the fly is in the center of the Y-Track. The flies are not able to determine which arm is rewarded simply by smelling or seeing reward. In trials where the fly turns in the non-reinforced direction, the servomotor is actuated rapidly so that the fly is never able to actually obtain food.

In order to validate the sensitivity of the real-time tracking, we compared long-term recordings of living flies and dead flies (*i.e*. flies that have no genuine locomotion; *n* = 2 per group; Figure 1D). We found that the tracked position of the dead flies was contained within a radius of 1 pixel over several hours. Locomotor episodes were therefore defined as continuous sequences of frames in which the fly moved at least one pixel, with the requirement that the fly must exceed a speed of 2 pixels/frame (0.38 mm/s) for at least one frame. Flies frequently paused between locomotor episodes, sometimes for extended periods of time. We defined pauses of greater than 1 minute as “rest” (Figure 1E).

Throughout the prototyping process, we evaluated the effectiveness of our apparatus in shaping wild type (WT) fly behavior. In pilot experiments (*n* = 15, WT flies, mixed sex), we found a small training effect of making sucrose available contingent upon turn direction in the center of the Y-Track. Interestingly, change in turn direction preference was correlated with time spent resting during training (Pearson’s *R* = 0.50). In order to rigorously test the hypothesis that rest is correlated with learning in the Y-Track, we trained a large cohort of flies (*n* = 85 female, 87 male; Figure 1F-I). Of this cohort, 11 (7%) rested early in training (3 minutes or more in Block #1-3). An additional 16 (9%) had high rest late in training (9 minutes or more in Block #4-6) and were excluded because WT flies reduce food seeking behavior during high sleep times (Donelson et al., 2012). Flies that did not rest (139 of 172) showed no increase in turn preference in the direction of reward, indicating that they did not learn the task (One-sample bootstrap test; *p* = 0.894). However, consistent with our pilot results, flies with early rest (11 of 172) had a significantly increased likelihood of turning in the direction of reward (One-sample bootstrap test; *p* = 0.031; Figure 1G-I). These results indicate that WT flies learn a sucrose-rewarded operant contingency only when they rest in the first half of the training trials. Because of the behavioral importance of these rest-defined groups, we will refer to flies that rest early in training as “drowsy” flies, and flies with low rest as “restless” flies.

### Y-Track Geometry Significantly Affects Thigmotaxis and Spontaneous Alternation

Operant conditioning paradigms designed for flies can be confounded by sensory cues; when presented with both an operant contingency and a classical prediction cue, flies preferentially attend to the classical cue (Brembs and Plendl, 2008). No classical cues were intentionally introduced into the Y-Track, but *post-hoc* examination of locomotor behavior of the in the apparatus revealed strong thigmotaxis behavior (Figure 2A). This is consistent with the behavior of flies in open-arenas (Simon and Dickinson, 2010), but it is potentially problematic for the Y-Track task for three reasons. First, if flies maintain contact with the wall through the vertex of the Y-Track, turn direction is correlated with a unilateral touch stimulus, which may act as a classical predictor. Second, it is unclear where the “choice point” for choosing a turn direction is located – presumably at whatever track location the flies “attach” to one of walls. Third, thigmotaxis may contribute to spontaneous alternation (Lewis et al., 2017), another behavior typical of unmanipulated WT flies. Spontaneous alternation would not independently result in flies preferring the rewarded turn direction, but, in simulated behavior, spontaneous alternation magnifies small turn biases into large turn preference indices (Figure 2C). If the effect of early rest is to modulate spontaneous alternation, it may be that restless and drowsy flies have the same mild change in “true” turn bias and the difference in turn preference index is due to changes in alternation.

**Figure 2.**
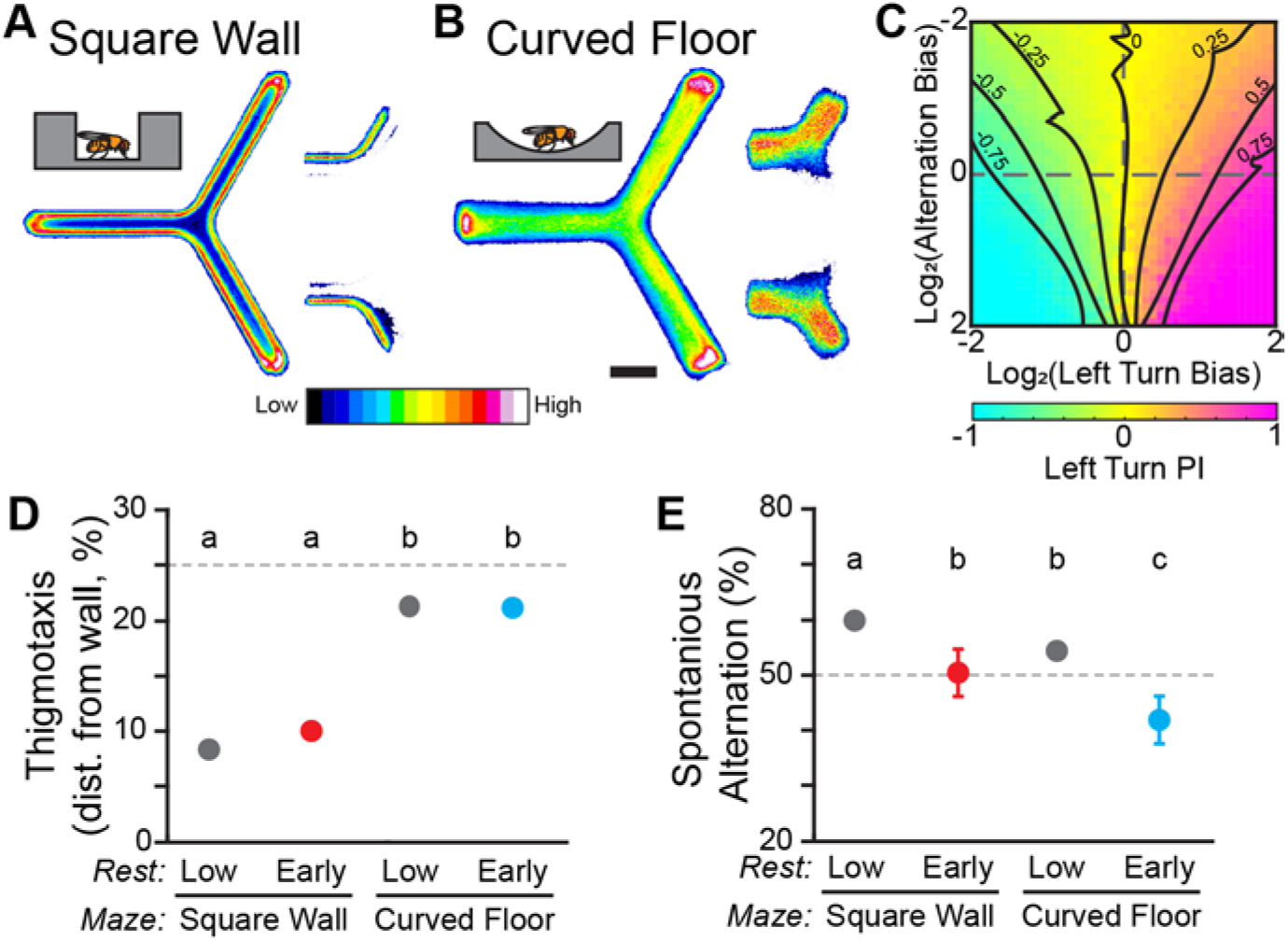
Y-Track Geometry Significantly Affects Thigmotaxis and Spontaneous Alternation. (A-B) Position heatmaps of flies in the square wall (A) and curved floor (B) Y-Mazes. Insets (*right*) show the heatmaps of trajectories through the center of the maze for flies walking from the left zone to the upper zone vs. the lower zone. Scale bar: 5 mm. Color shows relative occupancy of each pixel from low to high. (C) Heatmap of left turn Preference Index (PI) from a population of *in silico* flies. Each heatmap position is the mean of 1000 simulated trials of a fly with a range of left-turn bias and alternation bias. Dashed lines indicate zero bias on an axis. Solid lines are smoothed contours of constant PI. (D,E) Mean thigmotaxis (D) and spontaneous alternation (E) for WT flies in square wall and curved wall experiments. (D) Thigmotaxis is defined for each fly as the median distance from the wall of walking locations. The dashed line indicates the median distance from the wall if walking locations were distributed uniformly across the hallway. (E) Spontaneous alternation is measured in the final training block. The dashed line indicates the random rate of alternation. Error bars are standard error of the mean. Groups with the same letter are not significantly different from one another.

To address the sensory-motor confounds of thigmotaxis, we designed a second iteration of the Y-Track with gently curved floor, similar to open-field arenas (Simon and Dickinson, 2010). Thigmotaxis was dramatically reduced in the curved floor track compared to the square wall track, including as the flies pass through the vertex of the track (*n* = 103 WT female; Figure 2B). Median distance from the wall is significantly smaller in the square-wall design than it is in the curved floor design for both drowsy and restless flies (Two-way ANOVA; F(1,1) = 2788; *p* < 0.0001; Figure 2D). There was no difference in thigmotaxis between drowsy and restless flies within experiments (Post-hoc test; all *p* > 0.17).

To quantify any link between thigmotaxis, spontaneous alternation, and revealed turn preference, we measured spontaneous alternation at the end of training for each of our groups of WT flies. Track geometry significantly affected alternation rate (Two-way ANOVA; F(1,1) = 11; *p* = 0.001; Figure 2E) and drowsy flies have significantly lower alternation than restless flies (Post-hoc test; all *p* < 0.005). Although spontaneous alternation may be related to thigmotaxis and rest, spontaneous alternation changes in the wrong direction to confound the observed interaction between learning and rest.

### Learning to Turn Toward Sucrose is Independent of Track Geometry but Dependent upon an Informative Operant Contingency

The curved floor track geometry dramatically reduces thigmotaxis (Figure 2), so we repeated Y-Track conditioning in curved floor apparatus to determine if the sensory-motor experience of thigmotaxis contributes to Y-Track learning (Figure 3A). Learning in the curved floor track (*n* = 103 WT female flies) was similar to learning in the square wall track. The percentage of drowsy flies was not different between the square wall and curved floor tracks (Fisher’s Exact Test; *p* = 0.8). Drowsy flies in the curved floor track (8 of 103) significantly increased their likelihood of turning toward reward (One-sample bootstrap test; *p* < 0.0001) while restless flies (94 of 103) had no increase in turn preference (One-sample bootstrap test; *p* = 0.918; Figure 3B-D).

**Figure 3.**
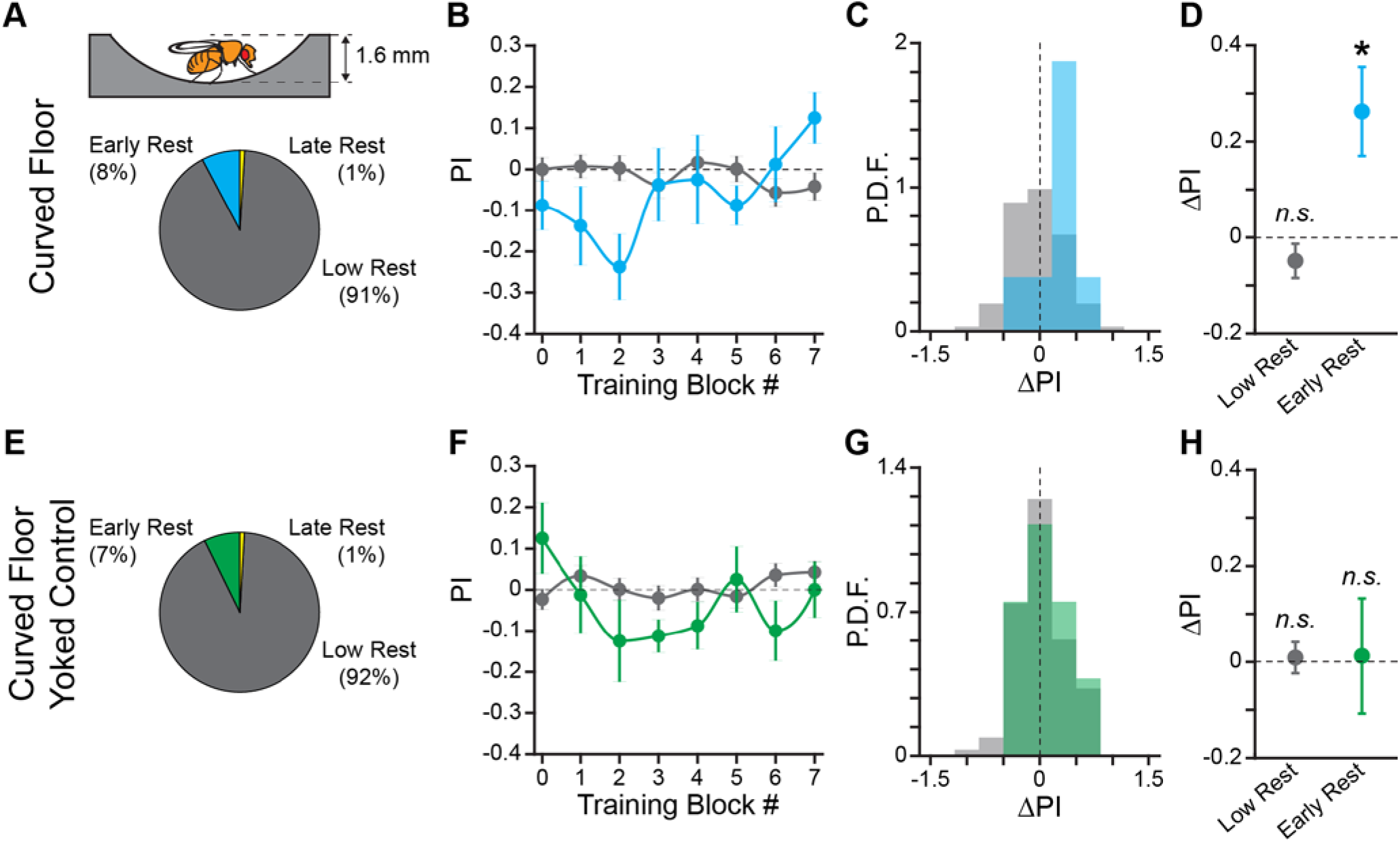
Learning to Turn Toward Sucrose is Independent of Track Geometry but Dependent upon an Informative Operant Contingency. (A) Schematic of hallway geometry with representative fly for scale (*Top*), Fraction of flies with each rest phenotype (*Bottom*). (B, F) Mean turn direction Preference Index (PI) for each training block for trained (B) and yoked control (F) flies in the curved floor Y-Maze. Points plotted in color are flies with early rest (drowsy), points plotted in grey are flies with low rest (restless). (C,G) Probability Distribution Function (P.D.F.) histograms of change in turn preference index (ΔPI) between training block #1 and #7 for trained (C) and yoked control (G) flies. Drowsy flies are plotted in color (*cyan*: trained, *green*: yoked), restless flies are plotted in grey. (D,H) Mean ΔPI of low rest (restless) and early rest (drowsy) flies trained in curved floor Y-Maze (D), and yoked control flies (H). Error bars are standard error of the mean, *n.s*.: groups are not significantly greater than zero, * groups are significantly greater than zero.

In order to control for any confounding effect of time spent in the experimental apparatus on turn direction preference, we also performed an open-loop, “yoked” control in the curved floor track (*n* = 110 WT female flies). In the yoked control flies, the reward sequence of a previously trained fly was presented to a naïve fly independent of the turning behaviors of the naïve fly. The yoked control flies therefore had the same amount of sucrose/reward access as trained flies, but there was no behavioral contingency. As predicted, neither drowsy (8 of 110) nor restless (101 of 110) yoked control flies significantly increase turn preference in the pseudo-rewarded direction (One-sample bootstrap test; *p* = 0.476 and *p* = 0.392, respectively; Figure 3E-H). Together, these results allow us to conclude that the change in turn preference is due to learning of the operant contingency and is not dependent upon sensory-motor feedback from thigmotaxis. The curved floor Y-Track was used for all subsequent experiments because learning is equivalent to the square wall track, without the potential confounding effect of thigmotaxis.

### A World Orientation Cue does not Facilitate Learning the Operant Task

The curved floor Y-Track experiment (Figure 3) eliminated the potential for an egocentric sensory-motor confounding cue. We next considered the possibility that the flies were attending to an inadvertently introduced world-orientation cue rather than the operant contingency. In order to test the hypothesis that flies are able to use orientation information to learn the task, we trained flies in the presence of a strong orientation cue (*n* = 81 WT female flies). A green LED was illuminated beneath one arm of the Y-Track, creating a stable, mildly attractive, landmark (Figure 4A). In the presence of this landmark, neither drowsy (9 of 81) nor restless (66 of 81) flies significantly increase turn preference in the rewarded direction (One-sample bootstrap test; *p* = 0.500 and *p* = 0.695, respectively; Figure 4B-D). This result indicates that the presence of a world orientation cue does not enhance, and may inhibit, learned change in turn preference. Together, the results of the thigmotaxis, alternation, and orientation-cue experiments indicate that neither sensory inputs nor motor patterns explain the change in turn preference in the direction of reward observed in drowsy flies. We conclude that the learning produced in this paradigm is navigational and operant in character.

**Figure 4.**
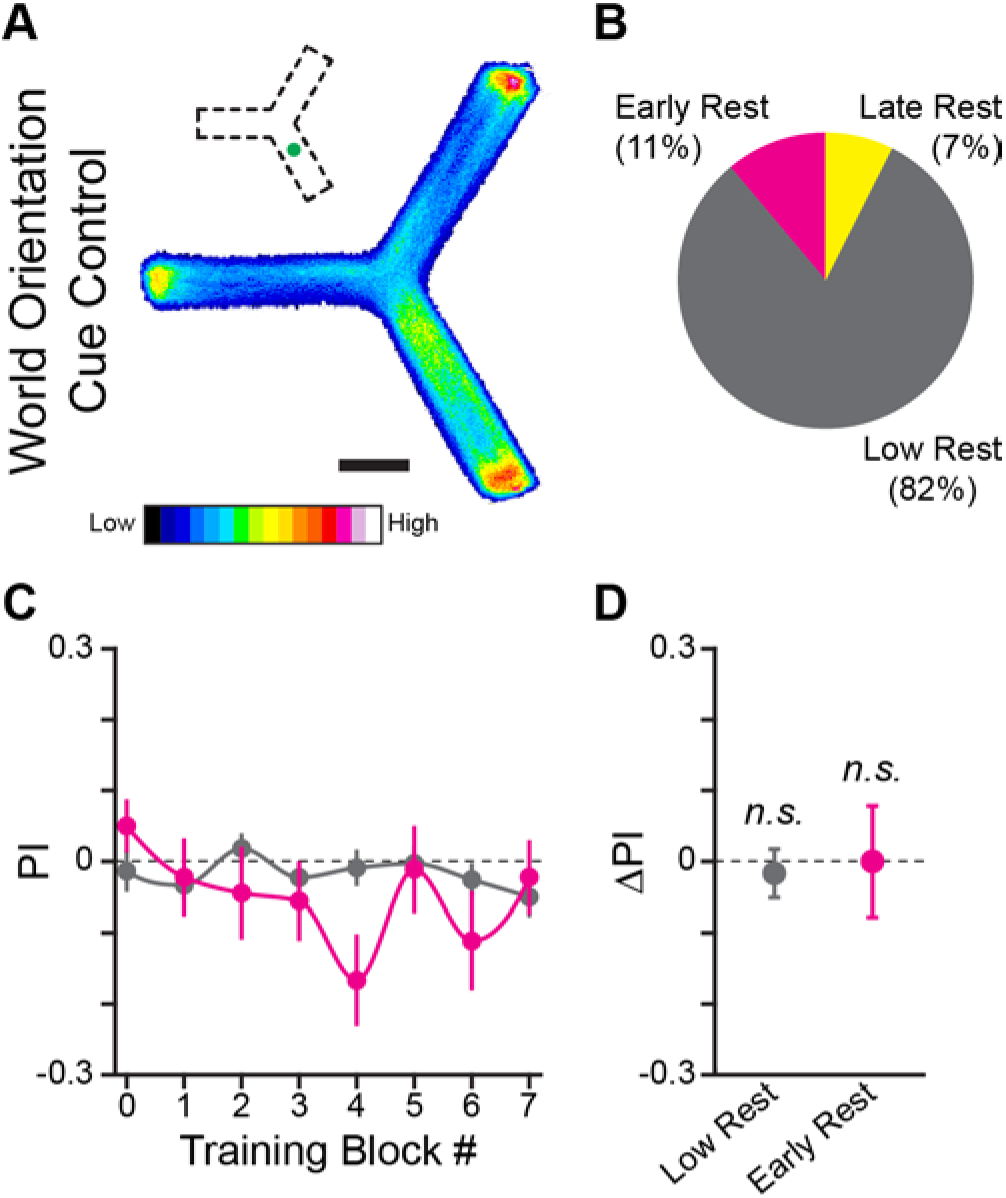
A World Orientation Cue does not Facilitate Learning the Operant Task. (A) Position heatmaps of flies trained with an LED spatial orientation cue. Scale bar is 5 mm. Color shows relative occupancy of each pixel from low to high. Small inset indicates the position of the green light cue. (B) Percentage flies with each rest phenotype. (C) PI by training block for early rest (magenta) and low rest (grey) flies. (D) Mean ΔPI of early rest and low rest flies. Error bars are standard error of the mean, *n.s*.: groups are not significantly greater than zero, * groups are significantly greater than zero.

### Sucrose Promotes Rest and is the Learning-Relevant US in the Y-Track

Operant conditioning is a learned association between behavior and a US, and the strength of the US influences the strength of the learned association (Rickard et al., 2009). Rest is strongly regulated by feeding and nutritional state (Murphy et al., 2016), and the consumption and hedonic value of sugar is dependent upon the state of the fly (Krashes et al., 2009; Li et al., 2020). While we have shown that there is a correlation between early rest and learning, it is not clear how they are connected mechanistically.

One possibility is that there may be a between-flies difference in the consumption of the sucrose US that explains enhanced learning in drowsy flies. In order to test the hypothesis that drowsy flies eat more sucrose and therefore receive a larger reward, we analyzed the relationship between time spent adjacent to sucrose and learning in the WT fly data reported above (Figures 1 and 3). We found that there is a significant main effect of early rest on time adjacent to sucrose where drowsy flies spend more time next to the sucrose (Two-way ANOVA; both F(1,1) > 4.5; both *p* < 0.035; Figure 5A). Importantly, both drowsy and restless flies spend enough time adjacent to the sucrose US (>5 seconds on average) to consume a sugar meal (approximately 2 seconds; Ro et al., 2014). Given this observed difference, we hypothesized that longer interactions with the US would result in greater learning. However, we found no correlation between time spent adjacent to sucrose and learning in the square-wall or curved-floor experiments (Pearson correlation; Square Wall *R* = 0.12, Curved Floor *R* = −0.01; Figure 5B). This analysis indicates that increased sucrose consumption is associated with additional rest but is not associated with increased learning.

**Figure 5.**
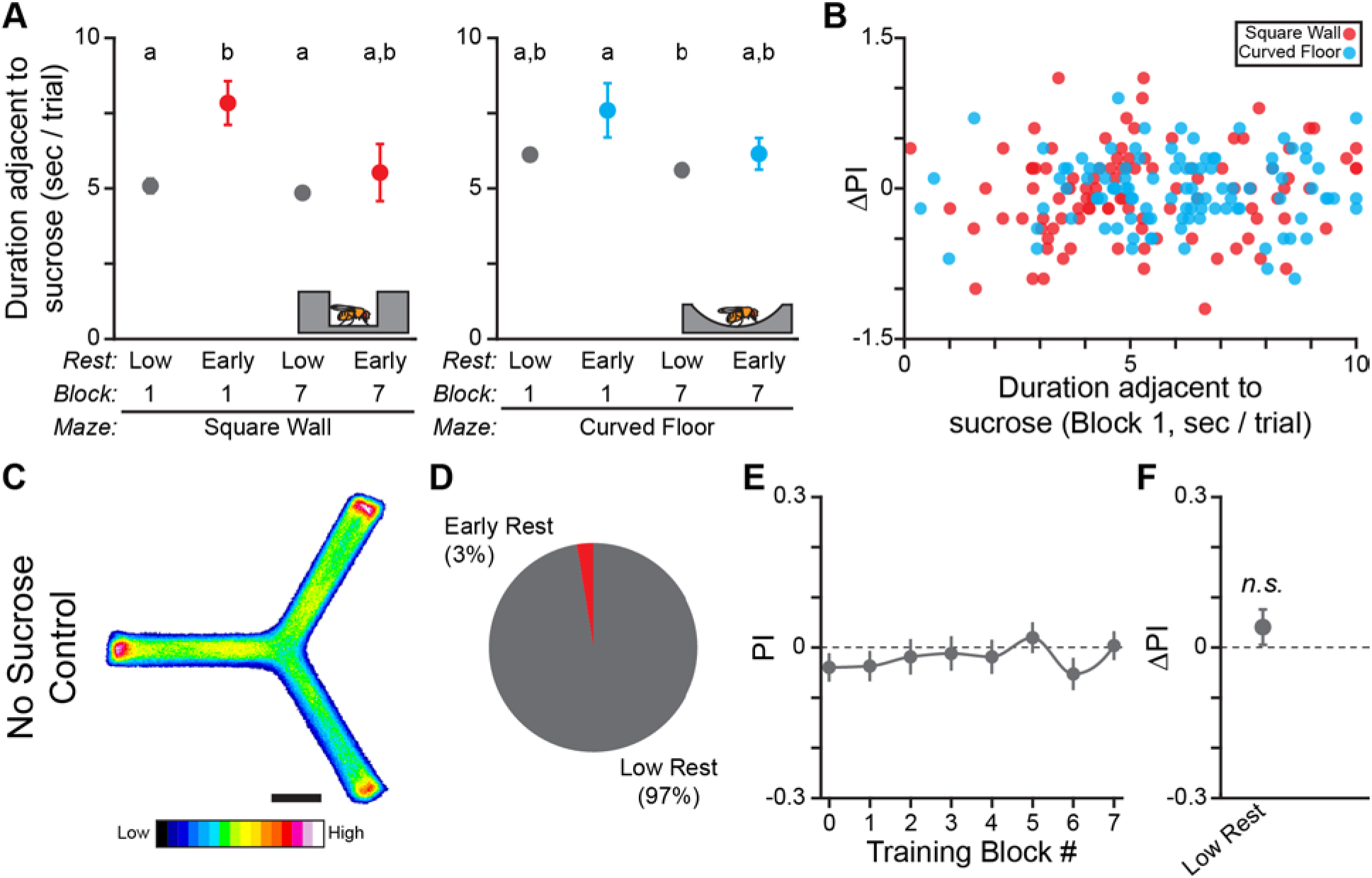
Sucrose Promotes Rest and is the Learning-Relevant US in the Y-Track. (A) Mean time per trial WT flies spend adjacent to sucrose in the square wall and curved floor experiments. Time per trial is capped at 10 seconds, because sucrose is automatically removed 10 seconds after it is made available. Groups with the same letter are not significantly different from one another. (B) Scatter plot of change in turn preference index (ΔPI) between training block #1 and #7 by time adjacent to sucrose in training block #1 for flies in the square wall experiment (red dots) and curved floor experiment (cyan dots). (C-F) Results of Y-Maze training with no sucrose available. (C) Position heatmaps of trained flies. Scale bar is 5 mm. Color shows relative occupancy of each pixel from low to high. (D) Percentage flies with each rest phenotype. PI by training block (E) and mean ΔPI (F) for low rest flies. Only 2 flies had early rest, which is insufficient to calculate a standard error, so these flies are not plotted. Error bars are standard error of the mean, *n.s.:* groups are not significantly greater than zero.

To validate this *post-hoc* analysis demonstrating that sucrose consumption promotes rest, we trained flies in a Y-Track with no sucrose available (*n* = 78 WT female flies). Remarkably, only 2 flies in this cohort had early rest, a 3-fold reduction in the fraction of drowsy flies compared to training with sucrose available (Figure 5D) and consistent with the sleep-suppressing effect of food deprivation on flies (Keene et al., 2010). The restless flies in the no-sucrose experiment did not have a significant increase in turn preference following training (One-sample bootstrap test; *p* = 0.119; Figure 5E-F). Together, increased time adjacent to sucrose in drowsy flies and the reduction in the number of drowsy flies when sucrose is removed from the Y-Track, indicate that sugar consumption promotes rest. The lack of residual learning of a secondary reinforcer in the Y-Track when sucrose is removed also shows that sucrose is the learning-relevant US.

### Optogenetically-Induced Sleep is Not Sufficient to Enhance Learning

Activation of several genetically-targetable cell types in flies is sufficient to induce sleep (Donlea et al., 2011; Liu et al., 2016). Activation of these cells has the same effect on sleep-dependent learning as spontaneous or pharmacologically-induced sleep (Donlea et al., 2011; Dissel et al., 2015). We therefore tested the hypothesis that sleep is sufficient to enhance learning of turn direction in the Y-Track by optogenetically activating sleep-promoting neurons. Flies do not synthesize the cofactor of Channel Rhodopsin (ChR), All-Trans Retinal (ATR), so ATR needs to be added to the food to functionalize the channels (Zhang et al., 2006). We tested optogenetic sleep induction using a dorsal Fan-Shaped Body driver (dFSB; 104y-Gal4; *n* = 39 ATR, 45 Vehicle), or an Ellipsoid Body driver (EB; VT058968-Gal4; *n* = 56 ATR, 54 Vehicle) driving expression of ChR2.XXL (Dawydow et al., 2014), and WT control flies (*n* = 43 ATR, 36 Vehicle) (Figure 6A). Sleep was induced by turning on blue LEDs located under the Y-Track for 5 minutes at the mid-point of Training Block 2 (Figure 6B). Lights were symmetrically located in all arms of the Y-Track, so they did not provide a world orientation cue. Blue light did not increase rest in WT flies fed ATR or vehicle, but dramatically increased rest in the dFSB and EB driver flies (Figure 6C,D). The drivers differed in both potency and on/off kinetics: the dFSB driver rapidly induces sleep in all ATR fed flies, and flies woke from sleep shortly after the light was turned off, consistent with prior reports (Tainton-Heap et al., 2020). In contrast, the EB driver induced a lower level of sleep that persisted even after the blue light was removed (Figure 6B). Both drivers dramatically increased the % flies with early rest (Figure 6D). Turn preference for reward was increased in drowsy WT flies (9 of 74), but not restless WT flies (62 of 74) (One-sample bootstrap test; *p* = 0.041 and *p* = 0.494, respectively; Figure 6E). Significant learning in the drowsy WT flies indicates that the changes in food and blue light illumination in this experiment did not prevent rest-dependent learning. However, turn preference for reward was not increased in either drowsy (43 of 84) or restless (36 of 84) flies in the dFSB induction experiment (One-sample bootstrap test; *p* = 0.775 and *p* = 0.964, respectively; Figure 6F). Nor was turn preference for reward increased in either drowsy (41 of 104) or restless (57 of 104) flies in the EB induction experiment (One-sample bootstrap test; *p* = 0.098 and *p* = 0.335, respectively; Figure 6G). The failure of drowsy flies to learn in the rest induction experiments implies that optogenetically-induced rest does not enhance learning. And because rest is induced in a large fraction of the flies without also enhancing their learning, the group of spontaneously drowsy flies that have learned is diluted with non-learning flies, effectively erasing the difference between the restless and drowsy groups. We conclude that spontaneous early rest is associated with increased learning of the navigational task, but that rest itself is not sufficient to induce learning.

**Figure 6.**
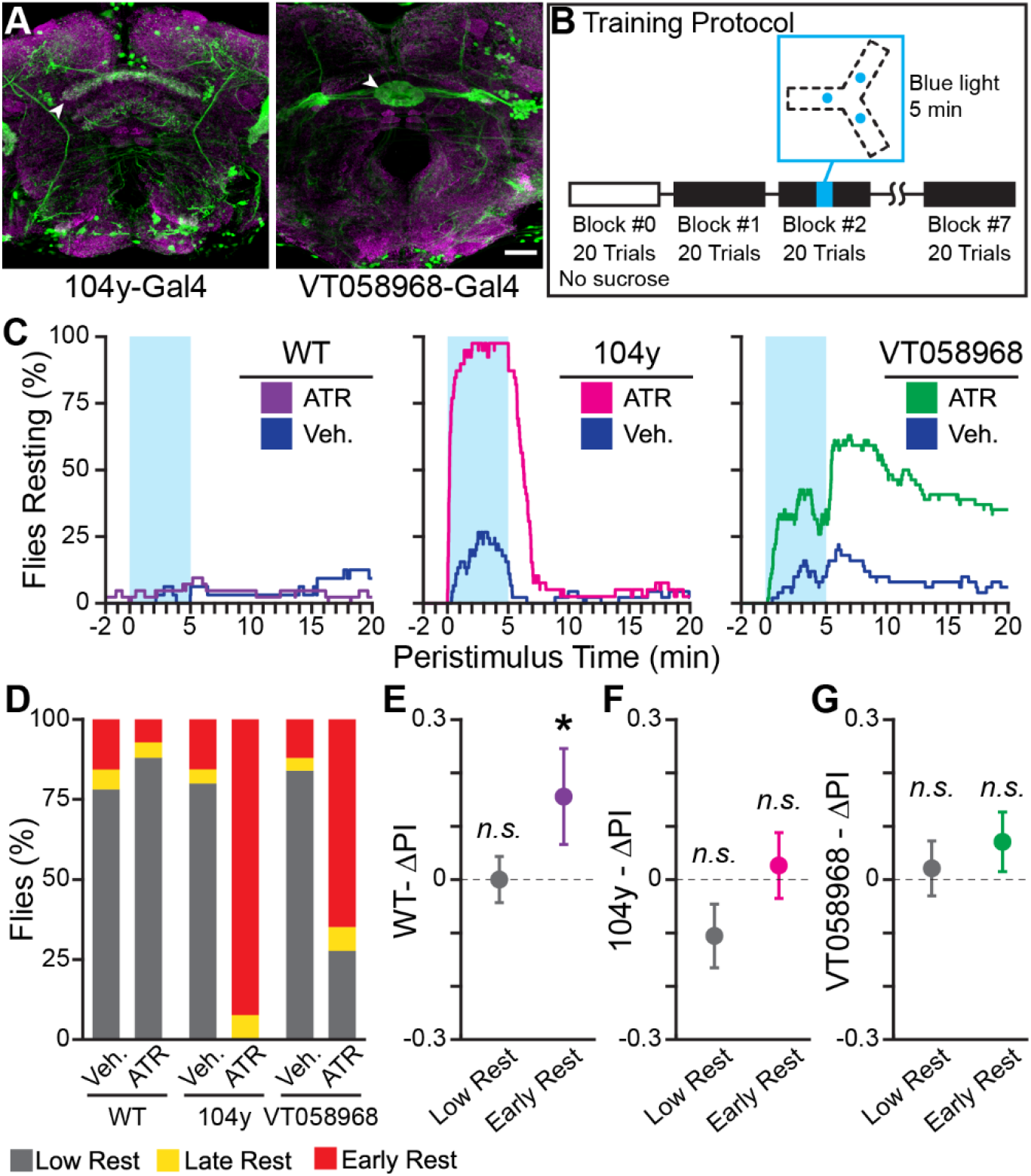
Optogenetically-Induced Rest is Not Sufficient to Enhance Learning. (A) Confocal maximum intensity projections of fly brains expressing 104y-Gal4;UAS-GFP (*left*) and VT058968-Gal4;UAS-GFP (*right*) stained for GFP (*green*) and Bruchpilot (nc82, *magenta*). Scale bar is 20 μm. White arrowheads indicate the sleep-promoting neuropil for each Gal4 driver: the dorsal Fan Shaped Body for 104y-Gal4 and the Ellipsoid Body for VT058968-Gal4. (B) Block diagram of operant conditioning paradigm, with the time of optogenetic stimulus indicated in cyan. The track diagram indicates the locations of each blue LED in the track, these positions are equidistant from the center of the track and illuminate the entire Y-track. (C) Peri-stimulus rest plots for WT (*left*), 104y (*center*), and VT058968 (*right*). Blue shaded area shows when the blue LED is on. The plots show 20 minutes of behavior following light onset - this time is not linked to trial times, which vary between animals, and does not encompass the entire duration of the experiment. (D) Percent of each rest phenotype present in each experiment group. (E-G) Results of training WT (E), 104y (F), and VT058968 (G) flies. Early and low rest flies for each genotype were pooled from the ATR and vehicle groups. Change in PI (ΔPI) is plotted for each genotype; colorful points are flies with early rest (*purple*: WT, *magenta*: 104y, *green*: VT058968). Error bars are standard error of the mean, *n.s*.: groups are not significantly greater than zero, * groups are significantly greater than zero.

### Y-Track Operant Learning is Dependent on cAMP Regulation

The Y-Track task we have developed is an operant, navigational, sucrose-reinforced learning paradigm, in which learning is revealed by changes in turn preference from baseline (Figure 1,3). Many neurotransmitters, receptors, and second messengers necessary for classical conditioning in flies have been identified using genetic knockouts (Margulies et al., 2005). While cyclic adenosine monophosphate (cAMP) is important for the formation of classical conditioning (Zars et al., 2000), a previous study found that activity-regulated cAMP synthesis is not necessary for the formation of aversive operant conditioning (Brembs and Plendl, 2008). In order to determine if regulation of cAMP is necessary for formation of appetitive operant conditioning, we tested the performance of flies mutant for the *dunce* phosphodiesterase (*dnc^1^*). *dnc^1^* flies fail to learn in classical paradigms and fail to modulate cAMP in response to learning stimuli (Gervasi et al., 2010). In an independent cohort (*n* = 209 WT female flies), we found that drowsy flies (19 of 209) have significantly greater learning than restless flies (183 of 209) (Two-sample bootstrap test; *p* = 0.035; Figure 7A-C). In contrast, flies carrying the *dnc^1^* mutation failed to show learning (*n* = 95 female flies): neither drowsy (11 of 95) nor restless (79 of 95) *dnc^1^* files significantly increased their likelihood of turning toward reward (One-sample bootstrap test; *p* = 0.181 and *p* = 0.591, respectively; Figure 7D-F).

**Figure 7.**
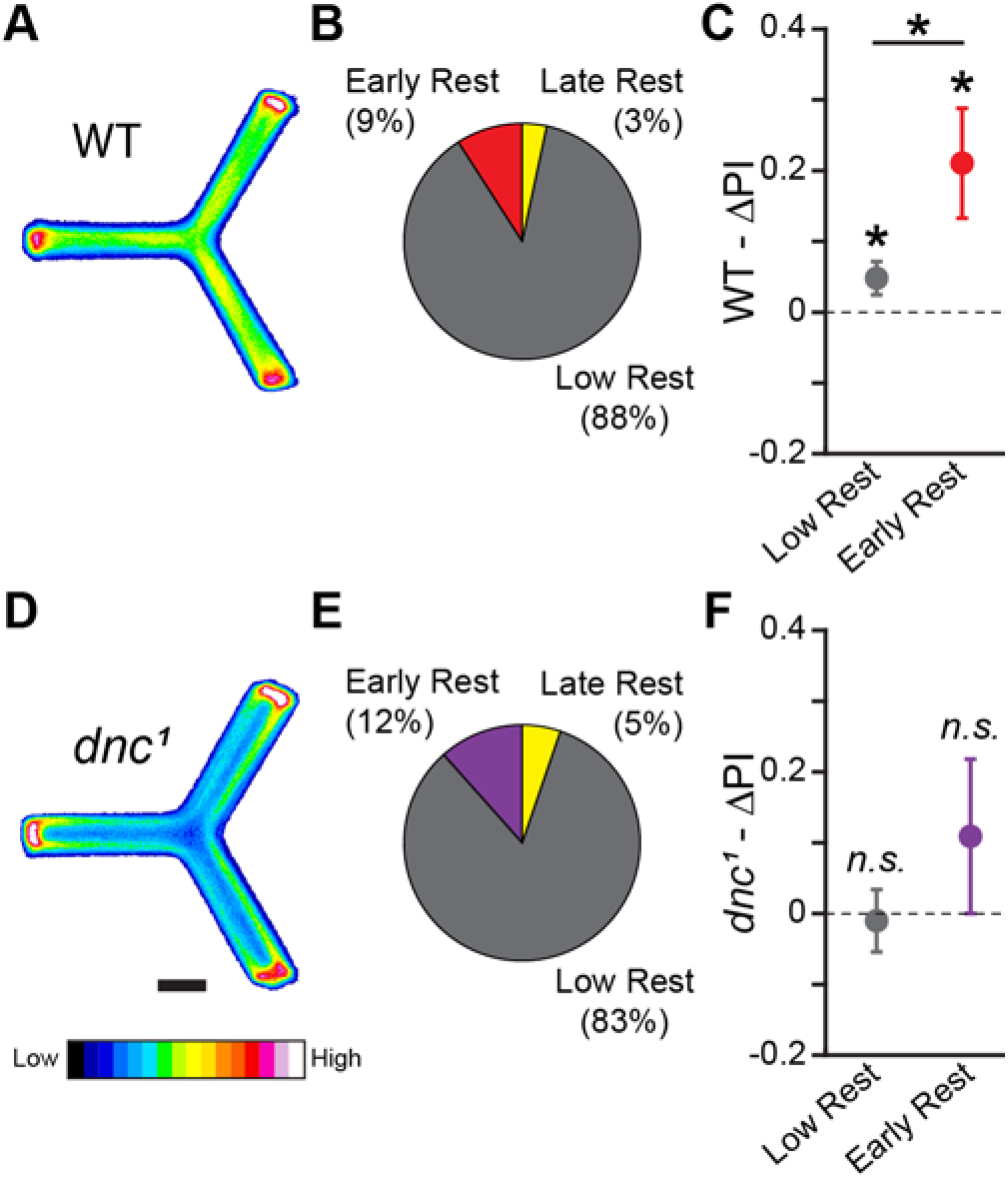
Y-Track Operant Learning is Dependent on cAMP Regulation. (A,D) Position heatmaps of trained WT (A) and *dnc^1^* (D) flies. Scale bar is 5 mm. Color shows relative occupancy of each pixel from low to high. (B,E) Fraction of WT (B) and *dnc^1^* (E) flies with each rest phenotype. (C,F) Mean ΔPI of low rest (restless) and of early rest (drowsy) WT (C) and *dnc^1^* (F) flies. The mean ΔPI of drowsy and restless WT files are both greater than zero (One-sample bootstrap test; *p* = 0.005 and *p* = 0.019, respectively). The mean ΔPI of drowsy WT flies is greater than the mean ΔPI of restless WT files (Two-sample bootstrap test; p = 0.035). Error bars are standard error of the mean, *n.s*.: groups are not significantly greater than zero, * groups are significantly greater than zero or different from one another.

## DISCUSSION

The formal study of associative learning has been remarkably successful: experimentally-induced associative memory has been demonstrated across the animal kingdom and dozens of genes, neurotransmitters, second messengers, and neural structures have been implicated in its formation (Mayford et al., 2012). Within the context of fly learning, animals have been trained to texture (Platt et al., 1980), sound (Menda et al., 2011), color (Schnaitmann et al., 2010), location (Wustmann et al., 1996), and odor (Quinn et al., 1974), among other cues. The fly learning literature developed rapidly, with first reports of training paradigms for classical conditioning to an aversive US, operant conditioning to an aversive US, and classical conditioning to a rewarding US occurring within a decade of one another (Quinn et al., 1974; Booker and Quinn, 1981; Tempel et al., 1983). In this report, we present the first paradigm for operant conditioning of flies to a sucrose US: Y-Track conditioning. The magnitude of the learning observed in WT flies (mean ΔPI = 0.2-0.3; Figure 1,3,7) is modest compared with immediate memory in highly-optimized olfactory learning paradigms (Flyer-Adams et al., 2020), but is similar to the magnitude of learning observed in early reports of olfactory and visual classical conditioning paradigms (Tempel et al., 1983; Vogt et al., 2014), and is also similar in strength to consolidated olfactory memories (Krashes et al., 2007). Like consolidated memory (Berry et al., 2015; Chouhan et al., 2020), the learning we observe is dependent upon rest (Figure 1,3). However, rest itself is not sufficient for learning as we see no learning enhancement in optogenetically induced rest (Figure 6), indicating that rest does not indiscriminately promote learning in the Y-Track. Y-Track learning does not depend on sensory cues (Figure 2,4) and requires cAMP as a second messenger (Figure 7). The properties of Y-Track learning have implications for navigation, learning and memory, and the connection between sleep and learning in *Drosophila*.

### Learning, Memory and Rest in Drosophila

Disorders of sleep and associative learning co-occur in several categories of neurological disease. Primary sleep disorders and sleep deprivation result in decreased cognitive and memory performance (Kessler et al., 2011; Shekleton et al., 2014; Zamarian et al., 2015). Conversely, plasticity disorders such as neurodevelopmental disability and post-traumatic stress disorder have co-morbid sleep abnormalities (Angriman et al., 2015; Gilbert et al., 2015). Finally, neurodegenerative disorders, including dementia, Huntington’s disease, and Parkinson’s disease, frequently disrupt both sleep and memory (Chaudhuri and Naidu, 2008; Morton, 2013; Robbins and Cools, 2014; Porter et al., 2015). The widespread connections between learning and sleep in human neurological disease indicate that there are neuronal circuits linking, or shared by, sleep and learning in humans. Similar to the sleep/learning connection in human disease, sleep and learning regulate one another in flies: sleep deprivation decreases learning (Seugnet et al., 2008; Chouhan et al., 2020) and learning increases time spent asleep (Ganguly-Fitzgerald et al., 2006). Remarkably, increased sleep is sufficient to rescue memory formation in learning mutant flies, aged flies, and in a fly model of Alzheimer’s disease (Donlea et al., 2014; Dissel et al., 2015), demonstrating that the shared circuit can be therapeutically useful.

Learning in the Y-Track conditioning task we have developed depends upon rest during training (Figure 1,3). We have described prolonged locomotor pauses as “rest” rather than “sleep” because sleep has a precise, three-fold definition (quiescence, increased arousal threshold, homeostasis) and it is not possible to properly evaluate this definition on the Y-Track (Hendricks et al., 2000; Shaw et al., 2000). We hypothesize that the locomotor pauses that we classify as rest are “sleep-like.” In light of the extensive links between sleep and learning in flies, we proposed two alternative hypotheses that could account for the correlation between rest and learning in the Y-Track: First, we hypothesized that learning could drive increased rest. This hypothesis is inconsistent with the similar amounts of rest observed in populations of flies that learn and those that do not (*i.e*. yoked controls and *dunce* mutants; Figure 3,7). Second, we hypothesized that rest drives increased learning. Sleep is known to promote consolidation of memory both in mammals and flies (Buhry et al., 2011; Donlea et al., 2011). However, we do not find that optogenetically-induced rest promotes learning (Figure 6). This failure to promote learning could be due to optogenetically-induced rest not promoting the learning-associated sleep state (Liu et al., 2019; Wiggin et al., 2020), as the brain-state of flies with dFSB-induced sleep is significantly different from the brain-state of spontaneous sleep (Tainton-Heap et al., 2020). It is also possible that the precise timing of rest is important to its learning-associated function and the timing of optogenetically-induced rest is suboptimal. Based on the failure of optogenetically-induced sleep to promote learning, we refine our hypothesis to propose that rest acts as a gate to learning. In this model, a coincidence between behavior and US must be detected, presumably by a cAMP-dependent mechanism (Figure 7). Following this coincidence detection event, sleep is required within a tight temporal window in order to consolidate the memory and prevent locomotion-related forgetting (Berry et al., 2015).

### Neural Circuits of Navigation as a Locus for Operant Plasticity

Identifying plastic neuronal circuits that are responsible for learning is a subject of intense interest in the effort to understand learning, memory, and cognition. In flies, the mushroom bodies are the best studied locus of learning-related plasticity. Learning of sensory cues in flies, including odors and visual cues, is mediated by mushroom body circuits (Aso et al., 2014; Vogt et al., 2014). The identification of the mushroom body as an important learning center proceeded primarily from neuroanatomy, including their connections to sensory projection neurons (Davis, 1993). Because operant conditioning is not primarily a sensory-driven behavior, a sensory-first search for neural circuits is unlikely to uncover the locus of plasticity that underlies operant learning. In fact, operant conditioning in the fly is mushroom body independent (Wolf et al., 1998; Brembs, 2009), while behavioral output circuits in the ventral nerve cord, such as motor neurons, have been implicated instead (Booker and Quinn, 1981; Colomb and Brembs, 2016). However, motor neurons themselves are unlikely to be the location of behavior/US coincidence detection (Talay et al., 2017). Motor planning circuits in the fly central complex, such as those responsible for navigation, are therefore an interesting potential neuronal locus for operant plasticity.

Control of turn direction on the Y-Track is determined by a mix of innate motor preferences and goal-directed search strategies. Innate handedness is strongly influenced by the activity of PB-FB-No neurons (PFN) (Buchanan et al., 2015). While the synaptic partners of PFN neurons implicated in innate handedness are not yet mapped, the brain structures innervated are all heavily involved in orientation and navigation in the fly (Giraldo et al., 2018; Shiozaki et al., 2020). Fly orientation circuits show rapid plasticity and features of short-term memory (Fisher et al., 2019) and are strongly responsive to visual stimuli (Seelig and Jayaraman, 2015). If fly orientation circuits are necessary for Y-Track operant conditioning, the mix of plasticity and visual responses would account for both behavioral plasticity and our finding that a strong visual stimulus inhibits learning rather than promoting it (Figure 4).

In addition to innate preferences, flies display strong learned place preference and goal directed search behaviors (Ofstad et al., 2011; Kim and Dickinson, 2017). Development of a location preference is capable of overriding innate preferences (Baggett et al., 2018) and foraging flies modify their innate locomotor preferences to repeatedly visit remembered sites of food and search for nearby food sources (Kim and Dickinson, 2017). The formation of spatial memories has been previously linked to cAMP as a coincidence detector (Zars et al., 2000), consistent with our finding that cAMP regulation is necessary for Y-Track conditioning (Figure 7). Our behavioral results are congruent with either plasticity happening directly in orientation/innate preference circuits, or in a foraging/place preference circuit. Further characterization of the neural components of spatial memory in flies is necessary for these possibilities to be distinguished.

## ACKNOWLEDGEMENTS

The authors would like to thank Hazal Uzunkaya and the Brandeis MakerLab for 3D printing assistance and would like to thank Junwei Yu for recommending the rest-promoting Ellipsoid Body driver (VT058968).

